# Artificial neural network language models predict human brain responses to language even after a developmentally realistic amount of training

**DOI:** 10.1101/2022.10.04.510681

**Authors:** Eghbal A. Hosseini, Martin Schrimpf, Yian Zhang, Samuel Bowman, Noga Zaslavsky, Evelina Fedorenko

**Author notes:** Corresponding authors: E. Hosseini, E. Fedorenko Conflict of interest statement: None.

## Abstract

Artificial neural networks have emerged as computationally plausible models of human language processing. A major criticism of these models is that the amount of training data they receive far exceeds that of humans during language learning. Here, we use two complementary approaches to ask how the models’ ability to capture human fMRI responses to sentences is affected by the amount of training data. First, we evaluate GPT-2 models trained on 1 million, 10 million, 100 million, or 1 billion words against an fMRI benchmark. We consider the 100-million-word model to be developmentally plausible in terms of the amount of training data given that this amount is similar to what children are estimated to be exposed to during the first 10 years of life. Second, we test the performance of a GPT-2 model trained on a 9-billion-token dataset to reach state-of-the-art next-word prediction performance on the human benchmark at different stages during training. Across both approaches, we find that (i) the models trained on a developmentally plausible amount of data already achieve near-maximal performance in capturing fMRI responses to sentences. Further, (ii) lower perplexity—a measure of next-word prediction performance—is associated with stronger alignment with human data, suggesting that models that have received enough training to achieve sufficiently high next-word prediction performance also acquire representations of sentences that are predictive of human fMRI responses. In tandem, these findings establish that although *some* training is necessary for the models’ predictive ability, a developmentally realistic amount of training (∼100 million words) may suffice.

## Introduction

A central objective in cognitive neuroscience is to develop models that can accurately predict human brain responses and behavior. In the neuroscience of language, some artificial neural network (ANN) language models were recently shown to be effective at predicting human brain activity and behavior during language processing (Caucheteux & King, 2022; Gauthier & Levy, 2019; Goldstein et al., 2022; Jain & Huth, 2018; Schrimpf et al., 2021; Toneva & Wehbe, 2019; Wilcox et al., 2020). For example, Schrimpf et al. (2021) examined the ability of over 40 language models to capture human responses to language and found that transformer architectures (Radford et al., 2019; Vaswani et al., 2017) fare best in aligning with human data. However, off-the-shelf models vary along many dimensions, making it difficult to unambiguously attribute any given model’s success in aligning with human data to particular model properties (architecture, objective function, amount/kind of training data, etc.). Gaining insights into human linguistic mechanisms requires controlled ‘experiments’ on the models, where different properties are systematically manipulated (Hu et al., 2020; Kumar et al., 2022; Warstadt & Bowman, 2019). This is the approach we adopt here in order to investigate how the *amount of training data* affects model-to-human alignment.

One common criticism of ANN models as models of human language processing is that their training data size (often, billions of words) far surpasses the amount of language exposure in humans during their learning phase (Chang & Bergen, 2021; Dupoux, 2018; M. C. Frank, 2023; Linzen & Leonard, 2018; van Schijndel et al., 2019); see (Warstadt & Bowman, 2022) for discussion). For example, (Hart & Risley, 1992) estimated that children are exposed to 3-11 million words each year, so by the time they turn 10 and possess adult-like linguistic competence, they are exposed to 30-110 million words. In contrast to a human child, who can learn a language from only ∼100 million words (or less), many current models get orders of magnitude more training data (20,000 human years’ worth for some models; Warstadt and Bowman 2022). More recently, (Gilkerson et al., 2017) estimated that by age of 10, the estimated amount of language exposure is around 20 million words, and (M. C. Frank, 2023) put this estimate at between 9 and 110 million words (extrapolating from their estimate of between 200 and 400 million by age 20). Here, we ask whether this extensive amount of training is necessary for the models to acquire representations that are predictive of human brain responses during language sentence comprehension.

Prior studies on the effects of training data on the models’ linguistic ability found that even with limited amounts of training data, models achieve considerable proficiency (Warstadt and Bowman, 2022). For example, Hu et al. (2020) and Zhang, Warstadt et al. (2021) report impressive syntactic generalizations in a BERT model (Devlin et al., 2018) trained on only millions of tokens (see also Huebner & Willits, 2021; Pannitto & Herbelot, 2020 for related evidence from a RoBERTa model trained on 5 million words of child-directed speech). And (Pérez-Mayos et al., 2021) find that a RoBERTa model (Liu et al., 2019) trained on 100 million words performs similarly to a model trained on 1 billion words on several syntactic benchmarks. These findings suggest that massive amounts of training may not be necessary for models to acquire certain aspects of linguistic competence. However, it is not known whether models trained on limited amounts of data can also predict human neural and behavioral responses to language.

Here, we evaluate how the *amount of training data* affects model-to-human alignment. In line with increasing emphasis in the field on robustness and replicability (Button et al., 2013; Ioannidis et al., 2014; Poldrack et al., 2017; Simmons et al., 2011), we adopt two complementary approaches (**Figure 1**). First, we investigate how well GPT-2 models (Radford et al., 2019) that are trained on different-size datasets (1, 10, 100 million, or 1 billion words) to reach their best training task performance, predict human fMRI and behavioral responses to sentences. Second, we investigate how a GPT-2 model’s ability to predict human fMRI and behavioral responses to sentences changes over the course of training on a large dataset in an effort to capture the ‘learning trajectory’ of model-to-brain alignment. In addition, we also examine the role of model perplexity in the ability of a model to predict human responses. To foreshadow the key results, we find that models reach high performance in predicting human responses to sentences even with realistic amounts of training data.

**Figure 1.**
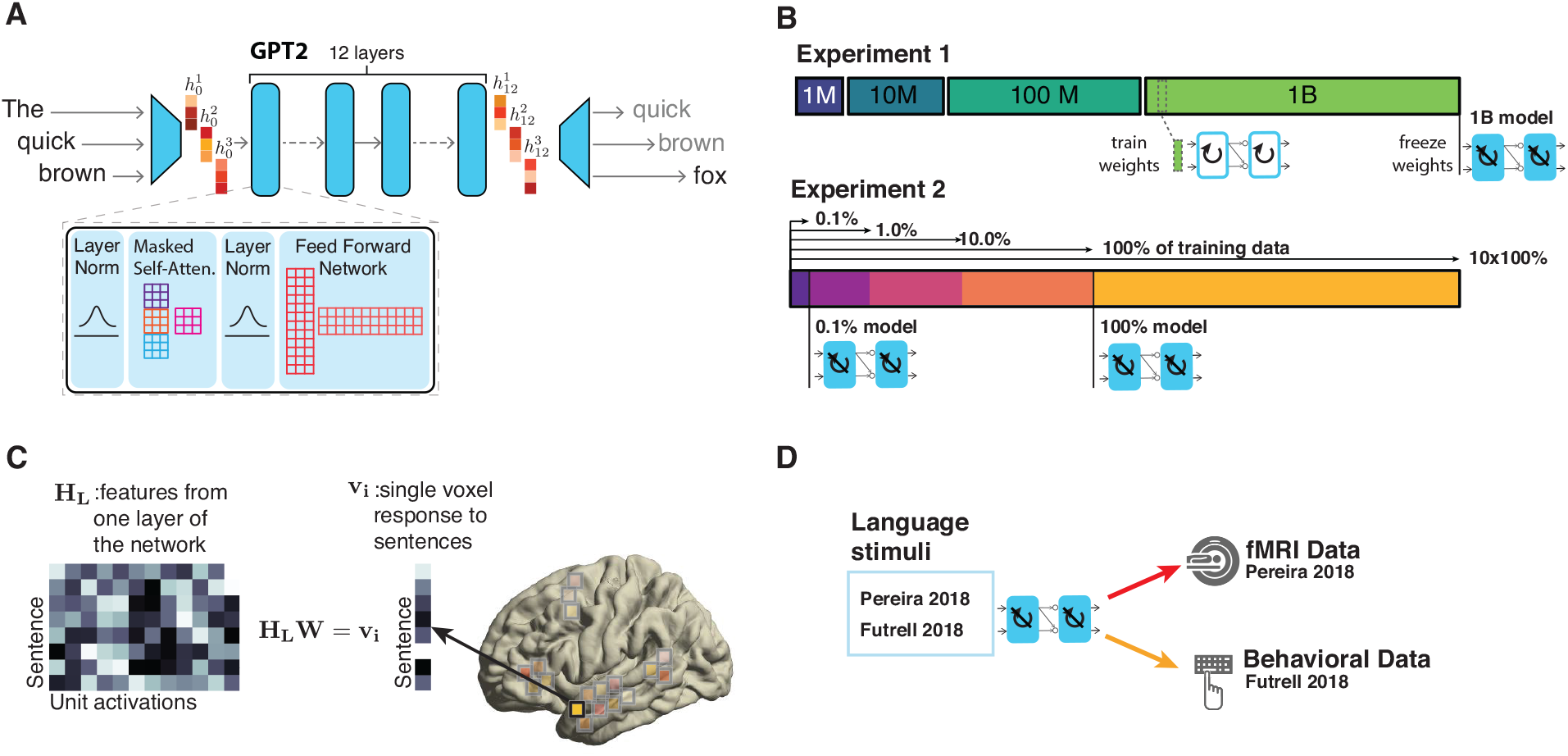
Methodological approach. **A.** Unidirectional-attention transformer architecture. Text input is processed sequentially to predict the next likely token at each step. **B.** The setup for Experiments 1 and 2. In Experiment 1, four models were trained using different-size datasets, and for each model, the weights with the best validation perplexity were frozen and used in the model-to-brain comparison. In Experiment 2, the GPT-2 model was trained using a very large dataset, and the weights were frozen at different steps during training and used in the model-to-brain comparison. **C.** Model representations were related to human representations by building a linear regression between unit activations for each layer of the

**Figure 2.**
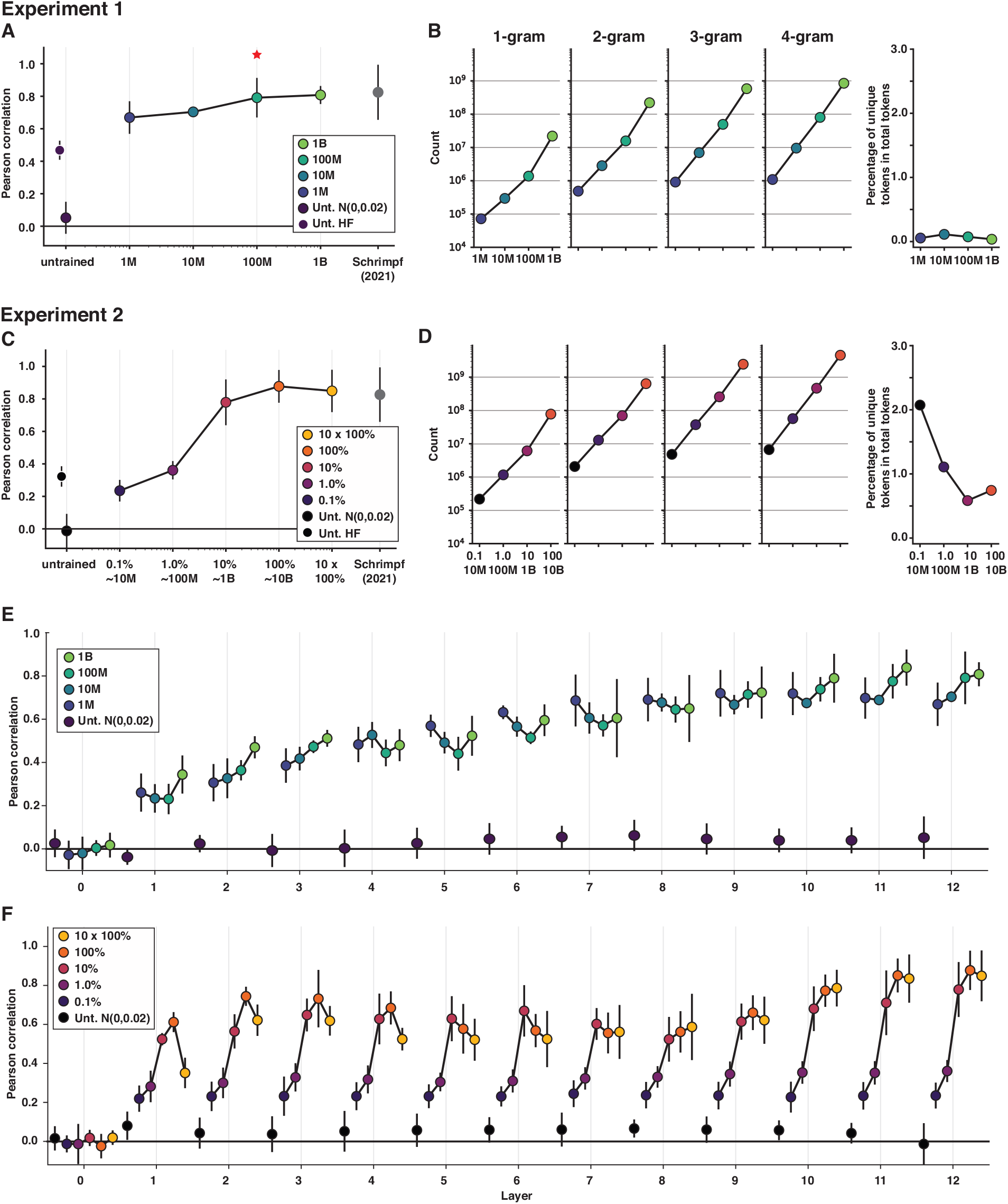
Model performance on the fMRI (Pereira2018) benchmark as a function of training. **A.** Experiment 1 results: performance (normalized predictivity; see text for information on raw predictivity) of the best-performing GPT-2 layer, as reported in Schrimpf et.al. (2021), in predicting language-responsive voxels’ activation in the Pereira2018 fMRI benchmark. The results are shown for i) two versions of an untrained (Unt.) model (initialized in two different ways: Unt. N(0,0.02) corresponds to the untrained model initialized with a mean of 0 and a standard deviation of 0.02, and Unt. HF corresponds to the untrained model initialized with the Hugging Face parameters; see <u>Methods</u>) (black dots); ii) four models trained on datasets of different size (1M, 10M, 100M, and 1B words) (blue-to-green dots connected by a line; the model trained on a developmentally plausible amount of data—100M—is marked with a red asterisk; see also **Supp. Figure X** for the results for a 50M model); and iii) a fully trained model, as reported in Schrimpf et al. (2021) (grey dots). Here, in (C), and in Figure 3, we computed a median score across participants and divided it by an estimated ceiling value to get a normalized score, and we computed a median absolute deviation over participants for use as error bars. **B.** The number of unique tokens (1-igrams), token 2-grams, token 3-grams, and token 4-grams in each training dataset. There is at least a logarithmic increase in the counts with the increase in the dataset size. The right-most panel shows the percentage of unique tokens relative to all tokens for each dataset. For Expriment 1, the total number of tokens is computed based on the number of training steps that was needed to reach best validation loss (see <u>Methods</u>; see also **Supp. Figure X** for the illustration of the training dynamics). **C.** Experiment 2 results: performance of the best-performing GPT-2 layer, as reported in Schrimpf et al. (2021), in predicting language-responsive voxels’ activation in the Pereira2018 fMRI benchmark. The results are shown for i) two versions of an untrained model (initialized in two different ways, as in A; see <u>Methods</u>) (black dots); ii) a model trained on a large dataset examined at different points during the training (0.1%, 1.0%, 10%, 100%, and 10*100% of training steps) (purple-to-yellow dots connected by a line); and iii) a fully trained model, as reported in Schrimpf et al. (2021) (grey dots). **D.** Same as in B but for the Openwebtext training dataset in Experiment 2. **E-F.** Exploratory analyses of individual model layers: performance of the 12 GPT-2 model layers in predicting human neural responses in the Pereira2018 fMRI benchmark in Experiment 1 (E) and Experiment 2 (F). (Layer 0 is the token embedding layer, and layer 12 is the last layer.) The results are shown for i) an untrained (Unt.) model (with the Gaussian initialization; black dots); and ii) four models trained on datasets of different size (blue-to-green dots connected by a line in E) or a model trained on a large dataset examined at different points during the training (purple-to-yellow dots connected by a line in F).

## Methods

### Experimental Design

#### Human datasets (benchmarks)

##### Primary benchmark: fMRI dataset (Pereira et al., 2018; referred to as “Pereira2018” below)

We used fMRI data from two experiments (Experiments 2 and 3 in Pereira et al., 2018). This benchmark is identical to the one reported in Schrimpf et al. (2021). Experiment 2 (n=9 native English speakers) consisted of 384 sentences across 96 Wikipedia-style passages, spanning 24 broad topics (professions, clothing, musical instruments, etc.). There were 4 passages per topic (for example, passages about a clarinet, an accordion, a piano, and a violin for the ‘musical instruments’ topic). Each passage consisted of 4 sentences, and the sentences varied in length between 7 and 18 words. Experiment 3 (n=9 native English speakers) consisted of 243 sentences across 72 passages, which were a mix of Wikipedia-style passages and short narratives. Each passage consisted of 3 or 4 sentences, and the sentences varied in length between 5 and 20 words. The stimuli for both experiments were constructed so as to span a broad range of topic areas. In both experiments, each sentence was presented on the screen for 4 seconds, followed by 4 seconds of fixation, and each participant read the materials 3 times across three fMRI scanning sessions. The responses were averaged across the three repetitions to derive a single response per sentence.

Furthermore, in each participant, the analysis was restricted to a set of voxels that were identified as language-responsive in an independent extensively validated language ‘localizer’ task (Fedorenko et al., 2010). In the localizer task, participants read sentences and list of nonwords (like ‘blork’ or ‘cre’) in a standard blocked design. Each item consisted of 12 words/nonwords that were presented one at a time at the rate of 450 ms per word/nonword.

Each sentence / nonword-list was followed by a simple button-press task (“press a button when you see a picture of a finger on a button”). Each trial lasted 6 seconds. Each block consisted of 3 trials and lasted 18 seconds. Each scanning runs consisted of 16 experimental blocks (8 per condition) and 6 fixation blocks and lasted 358 seconds. Each participant completed two runs, and the order of conditions was counterbalanced across runs. The localizer is available for download at (anonymous link). The contrast between sentences and nonword lists has been shown to robustly identify the fronto-temporal language-selective network of brain areas (Fedorenko et al., 2011; Lipkin et al., 2022b). These areas support language comprehension across modalities (listening, reading, etc.) and have been established to be sensitive to both word meanings and syntactic structure processing (e.g., Fedorenko et al., 2010; Fedorenko et al., 2020). For each participant, we selected the top 10% of most localizer-responsive voxels within a set of twelve broad masks (six in each hemisphere) that cover inferior frontal and lateral temporal cortex (the masks were derived from an independent set of 220 participants who performed the language localizer task and are available at (anonymous link). Thus, the fMRI benchmark consists of—for each of the two experiments—a set of language responsive voxels in each participant, and for each voxel, we have an estimate of the BOLD response to each of 384 sentences (Experiment 2 in Pereira et al., 2018) or 243 sentences (Experiment 3 in Pereira et al., 2018).

##### Secondary benchmark: Behavioral (reading-times) dataset (Futrell et al., 2018; referred to as “Futrell2018” below)

We used self-paced reading data from (Futrell et al., 2018). Similar to the fMRI benchmark, this benchmark is identical to the one reported in Schrimpf et al. (2021). 179 native adult English-speaking participants (recruited through Mechanical Turk) read stories that were presented one word at a time; with each button press, the current word would disappear in place of the new word (e.g., Just et al., 1982). The time it took a participant to move to the next word, n+1, was used as a measure of comprehension difficulty at word n. The stories were based on existing stories but were edited in a way so as to increase the frequency of rare words and constructions, including constructions that are known to cause comprehension difficulty (see Futrell et al., 2018, for details). The stories consisted of 33-64 sentences, and contained between 938 and 1,089 words. Each of 179 participants read between 5 and 10 stories and answered comprehension questions at the end of each story; each story was read by 82-98 participants. Following Futrell et al. (2018), we excluded reading times outside of the [100ms, 3,000ms] range. Note that we only report the results for this benchmark in **SI Figure 3** because, as will be discussed below, we found that diverse variants of control (untrained) language models predict these responses well above chance, which suggests that model predictivity is unlikely to be related to the representation of the linguistic stimuli.

#### Artificial Neural Network Models

We used two different implementations of a GPT-2-style model. For Experiment 1, where a model was trained on a dataset with a controlled number of words, we used the GPT-NEOX library which is a distributed training framework that uses the DeepSpeed library (Black et al., 2022; Aminabadi et al., 2022). We used a unidirectional-attention transformer model (GPT-2; Radford et al. 2019) with 12 layers and an embedding layer which was learned during training. Each layer had a size of 768 units and consisted of 4 main blocks (**Figure 1A**): (i) 1st layer normalization, (ii) self-attention, (iii) 2nd layer normalization, and (iv) the feedforward layer. The final layer consisted of a linear projection with a sigmoid nonlinearity that mapped hidden states into probabilities over the dictionary. The context size was 1,024 tokens. To test whether our results would generalize to bidirectional-attention transformer architectures, we additionally used publicly available miniBERTa models^1^ that were trained on the same datasets as the GPT-2 models (Zhang et al., 2020). (We did not include the model trained on the smallest (1 million words) dataset, for which Zhang et al. (2020) used a smaller-size model, which therefore would not be directly comparable to the other models.) The miniBERTas use the same design as the RoBERTa ‘base’ model (Liu et al., 2019)—a bidirectional-attention model with 12 layers, each 768 units in size, and a context size of 512 tokens. Importantly, RoBERTa has the same number of parameters as GPT-2 (125 million), allowing for a relatively controlled comparison of uni- and bidirectional architectures.

For Experiment 2, to investigate model training dynamics with a very large dataset, where during the early stages of the training the model continues to see new input (cf. doing multiple passes through a smaller-size training corpus as in Experiment 1), we used GPT-2 model weights from a publicly available model from the HuggingFace Transformers library (https://huggingface.co/stanford-crfm). The model has a similar architecture to the GPT-2 model used in Experiment 1.

#### Model Training

##### Training datasets

For Experiment 1, we combined the BookCorpus (Zhu et al., 2015) and English Wikipedia (Liu et al., 2019; Zhu et al., 2015) with a 1:3 ratio. We then created 4 different datasets with 1 million, 10 million, 100 million, and 1 billion words. These were used for training both the GPT-2 models and the minBERTa models. For Experiment 2, we used a model that was trained on the OpenWebText corpus (Gokaslan & Cohen, 2019) with more than 9 billion tokens.

To characterize the training corpora, we counted the number of unique tokens, token bi-grams, token tri-grams and token four-grams for different dataset sizes in Experiment 1 and for different checkpoints in Experiment 2. For this analysis, the data were tokenized as in the Penn Treebank corpus (Marcus et al., 1993). In particular, contractions were split (e.g., they’re è they+ re) and punctuation marks were treated as separate tokens. Afterwards, we counted unique occurrences of tokens, tokens bi-grams, and so on. As shown in **Figure 2B,D**, the number of all n-grams increases with corpus size (Experiment 1) and for later checkpoints (Experiment 2), and the percentage of unique tokens relative to total tokens is always higher in Experiment 2 compared to Experiment 1 (see also **Supp. Figure 10** for an illustration of the training dynamics in Experiment 1 vs. 2). (Quantifying the variability in syntactic structures is more challenging given the size of the corpora and the fact that they are not parsed/POS-tagged.)

In addition, following a reviewer’s request, we tested whether the experimental materials from the human benchmarks were present in the training corpora. We found that none of the sentences from the fMRI benchmark were present in any of the training corpora, and only a very small number of sentences from the behavioral benchmark were present in the training corpora. In particular, in Experiment 1, 3 of the sentences from the behavioral benchmark were found in the 1M training dataset, 2 sentences – in the 10M training dataset, 4 sentences were found in the 100M training dataset, and 17 sentences in the 1B training dataset; in Experiment 2, 4 sentences were found in the training dataset.

##### Training procedure

For Experiment 1, to train the GPT-2 models, we used standard initialization from the GPT-NEOX library and standard training parameters (Radford et al., 2019; see **Suppl. Figure 1** for details). After training, the model weights with the smallest validation perplexity were selected for evaluation on the human benchmarks. The smallest validation perplexity was reached after 1,000 steps for the 1M dataset (1,024 tokens * 128 batches * 1,000 steps = **131,072,000** tokens total), after 2,000 steps for the 10M dataset (1,024 tokens * 128 batches * 2,000 steps = **262,144,000** tokens), after 14,250 steps for the 100M dataset (1,024 tokens * 128 batches * 14,250 steps = **1,867,776,000** tokens), and after 310,000 steps for the 1B dataset (1,024 tokens * 128 batches * 310,000 steps = **60,948,480,000** tokens). To train the miniBERTa models, we used standard initialization and training parameters from the Hugging Face Transformers library (Liu et al., 2019).

Predictivity of the trained models was compared to that of untrained models. Here, in addition to the untrained GPT-2 model available from the Hugging Face library, we created an alternative untrained GPT-2 model in order to investigate the effects of different weight initializations on the alignment between model representations and human neural responses and reading behavior, and thus to isolate the effects of model architecture alone (i.e., the units and the patterns of connections among them) on predictivity. This version implemented the same unidirectional mask as the trained models and the other untrained model, but all the weights were set to a gaussian distribution with a fixed mean and standard deviation (mean: 0, standard deviation: 0.02 for the layer normalization, self-attention, and feedforward layer weights; see **Supp. Figure 6** for a detailed comparison with the Hugging Face initialization parameters).

For Experiment 2, the GPT-2 model was trained with standard initialization and training parameters until it reached state-of-the art perplexity values. We selected several checkpoints at which we extracted model representations from each layer for evaluation on the human benchmarks. The model was trained on 16 GPUs, with 8 batches per GPU, and updates were performed after 4 gradient accumulations. As a result, a *training step* constituted 1,024 (tokens) * 8 (batches) * 16 (GPUs) * 4 (gradient accumulations) = 524,288 tokens. Given that the tokenized OpenWebText corpus contains 9,036,044,288 tokens, it takes close to 20K training steps (specifically, 17.2K steps) to do one complete pass over the corpus. The checkpoints were selected in a logarithmic manner (0, 0.1% (20 training steps, which corresponds to 524,288 tokens * 20 steps = 10,485,760 tokens), 1.0% (200 steps, which corresponds to 524,288 tokens * 200 steps = 104,857,600 tokens), 10% (2K steps, which corresponds to 524,288 tokens * 2,000 steps = 1,048,576,000 tokens), 100% (20K steps, which corresponds to 524,288 tokens * 20,000 steps = 10,485,760,000 tokens), and 10×100% (200K steps, which corresponds to 524,288 tokens * 200,000 steps = 104,857,600,000 tokens)).

#### Analyses

##### Model comparison to the fMRI benchmark

We followed the approach in Schrimpf et al. (2021). In particular, we first extracted the representation for all the sentences that were used in the human fMRI experiments from each layer of each model. For each experiment, we split the stimuli into 5 ∼equal-size batches (Experiment 1: 3 batches of size 76 sentences and 2 batches of size 78; Experiment 2: 2 batches of size 48 sentences and 3 batches of size 49 sentences) and used ∼80% of the data to build an ordinary least squares regression model between model unit activations and voxel-level responses in the language network (defined by an extensively validated language ‘localizer’ task, as described in <u>Methods</u>; (Fedorenko et al., 2010; Lipkin et al., 2022a). For both experiments, we selected the representation of the last word in each sentence (for multi-token words, we averaged the representations across the composite tokens). We then applied the regression to the left-out ∼20% of sentences to generate predictions for BOLD responses in each voxel and compared these predictions against the observed BOLD responses using Pearson correlation. This procedure was iterated across the data folds, leaving out a different 20% each time, and the Pearson values were averaged across these iterations in each voxel of each participant. For each participant averaging across the two experiments (Experiments 2 and 3 in Pereira et al., 2018), we obtained a single score by taking the median Pearson value across the language-responsive voxels. A reliable positive Pearson correlation value indicates that the model is able to predict fMRI responses with a linear transformation. The resulting Pearson correlation values are divided by the ceiling value computed by estimating how well a best possible model of an average human would predict fMRI responses (Schrimpf et al., 2021). This value was estimated to be 0.32 for the Pereira et al. (2018) dataset. (We acknowledge that the issue of how to compute a noise ceiling remains an open issue in the field.)

Statistical testing was performed on the scores from the best model layer (as determined in Schrimpf et al., 2021) and took the form of independent two-sample t-tests. In particular, participant scores for each model in Experiment 1 (n=5 models: untrained, 1M, 10M, 100M, and 1B) or each checkpoint of a model in Experiment 2 (n=6 checkpoints: untrained, 0.1% of training steps, 1% of training steps, 10% of training steps, 100% of training steps, and 10×100% of training steps) were compared to the scores for the fully trained model (from Schrimpf et al., 2021). The resulting p-values were Bonferroni-corrected for the number of comparisons (5 and 6 comparisons in Experiments 1 and 2, respectively).

##### Model comparison to the behavioral benchmark

Similar to the fMRI benchmark, we followed the approach in Schrimpf et al. (2021). In particular, we first extracted the representation for all the stimuli (individual words) that were used in the behavioral experiment from the last layer of each model. For each participant, we split the total words (which varied across participants depending on the number of stories that a participant read) into 5 ∼equal-sized batches, and used ∼80% of the data to build an ordinary least squares regression model between model unit activations and reading times. In dividing the data into batches, we ensured that a) the same word did not appear in the training vs. the test set, and b) for any given sentence, the words were divided as evenly as possible between the training and the test set. We then applied the regression to the left-out ∼20% of words to generate predictions for reading times and compared these predictions against the observed reading times using Pearson correlation. This procedure was iterated across the data folds, leaving out a different 20% each time, and the Pearson values were averaged across these iterations to obtain a single score per participant. A reliable positive Pearson correlation value indicates that the model is able to predict reading times with a linear transformation. The resulting Pearson correlation values are divided by the ceiling value computed by estimating how well a best possible model of an average human would predict reading times (Schrimpf et al., 2021). This value was estimated to be 0.76 for the Pereira et al. (2018) dataset. Statistical testing was performed on the scores from the last model layer and took the form of independent two-sample t-tests, with a Bonferroni correction for the number of tests, similar to the fMRI benchmark.

##### Model perplexity

Following standard practice (e.g., Jelinek et al., 1977), we used perplexity as a measure of model performance on the language prediction tasks (next-word prediction for the GPT-2 models and missing-word prediction for the miniBERTa models). Perplexity (PPL) is defined as:

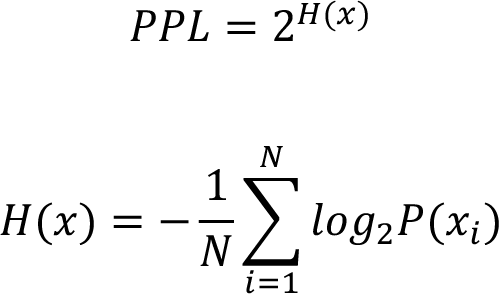

where H is entropy, and x denotes the tokens.

For both experiments, we used the test set from the wikitext-103-raw-v1 dataset (Merity et al., 2016) to compute perplexity. Perplexity was computed using a context size of 1,024 tokens and a stride of 512 tokens.

##### Code Accessibility

The human benchmarks and the code for relating model representations to the benchmarks are publicly available at https://github.com/mschrimpf/neural-nlp. For Experiment 1, the GPT-2 models and the representations extracted for all the benchmarks are available upon request (and will be uploaded to the Hugging Face library upon the paper’s publication); the miniBERTa models are available upon request; the checkpoints are available at https://huggingface.co/nyu-mll. (The training corpora used for Experiment 1 have copyright restrictions so cannot be made publicly available.) For Experiment 2, the model checkpoints are available at https://huggingface.co/stanford-crfm; the training corpus is available through the Hugging Face library.

## Results

### 1a. Models that are trained on relatively small amounts of data (small corpora) can predict human fMRI responses to sentences

We started by examining the performance of a unidirectional transformer model trained on the standard language modeling task in predicting human fMRI responses during the processing of sentences. Specifically, we tested a GPT-2 architecture (**Figure 1A**), which has previously been shown to best capture human neural and behavioral responses (Schrimpf et al., 2021). We trained four independent models on 1 million, 10 million, 100 million, and 1 billion words, respectively (**Suppl. Figure 1**). A training dataset of size 100 million words is comparable to the amount of language input that children have been estimated to get during the first decade of life (M. C. Frank, 2023; Hart & Risley, 1992). (Of course, the *nature* of the language input is still quite different between models and children both with respect to the content and the modality (text only for models vs. multi-modal input for children). We return to this point in the Discussion.) After training, we selected the checkpoint with the best perplexity on the validation set and tested how well the model representations capture human neural (fMRI) responses to sentences in the language-selective network (Fedorenko et al., 2011) and human behavioral responses in a self-paced reading task (**Figure 1D**).

For the primary, fMRI (Pereira2018) benchmark (Pereira et al., 2018; Schrimpf et al., 2021), we observed a consistent increase in performance with an increase in the size of the training set (**Figure 2A**; see **Suppl. Figure 5** for evidence—for this and the behavioral benchmark—of robustness of this pattern to seed choice during model initialization; cf. (Mehrer et al., 2021); see Frank et al., 2015 and Aurnhammer & Frank, 2020 for similar findings from earlier, pre-transformer models, including ngram, RNN, and phrase-structure grammar models). Critically, however, the model trained on just 100 million words already exhibits fMRI response predictivity that is similar to that of the fully trained GPT-2 model as reported in Schrimpf et al. (2021), with no significant difference in predictivity values (p=0.99; the data frames are available at OSF: anonymous link). The model trained on 1 billion words also does not differ from the fully trained model in predictivity (p=0.82). In contrast, the predictivity of the untrained model (the version with the Hugging Face initialization parameters) and the models trained on 1 million and 10 million words is significantly below the predictivity of the fully trained model (p-values: <0.0001, 0.007, and 0.016, respectively; here and elsewhere, the values are Bonferroni-corrected, as described in Analyses).

The untrained model performance differs between the two versions (see <u>Methods – Training Procedure</u>). The version initialized with the standard Hugging Face parameters performs well above chance (p<0.0001), as reported in Schrimpf et al. (2021) (see also (Caucheteux & King, 2022), but the version initialized with the alternative parameters (all weights set to a normal distribution with a mean of 0 and a standard deviation of 0.02) performs around 0 (not significantly different from 0; p=0.14) (**Figure 2A**).

The results also generalize, to some degree, to a bidirectional transformer model (miniBERTa; Liu et al. 2019) (**Supp. Figure 2**). In particular, similar to the GPT-2 models, we observed a consistent increase in model performance with an increase in the training dataset size, which suggests that this pattern is robust to architecture. However, the 100M word model still performs below the fully trained model. This difference between the GPT-2 and miniBERTa models in the amount of training they require to align with human data is likely due to the difference in the directionality of the attention mechanisms, with unidirectional-attention mechanisms being more sample-efficient. Generalizing these results to other minimally different variants of uni-vs. bidirectional-attention transformer models will help strengthen this conclusion.

In exploratory analyses, in addition to examining the language network as an integrated system, we examined the effects of the amount of training data on the models’ ability to predict fMRI responses in individual frontal and temporal language fROIs (for a total of six language fROIs in the left hemisphere, and six homotopic regions in the right hemisphere; e.g., Lipkin et al., 2022). The results are shown in **Supp. Figure 9**. The overall pattern was similar across all language fROIs, including between the LH IFG fROI and the LH PostTemp fROI (which have been argued by some to differ functionally; e.g., (Friederici, 2018; Hagoort, 2019); cf.(Fedorenko et al., 2020). The overall predictivity was lower in the RH than the LH language fROIs (p <<0.0001 for all models in Experiment 1 and all checkpoints in Experiment 2), in line with past findings (e.g. (Schrimpf et al., 2021; Tuckute et al., 2023)).

We also investigated the patterns of model-to-brain alignment across model layers. Prior work in vision (Geiger et al., 2020; Storrs et al., 2021) has suggested that training affects model performance differently across layers, with early layers already reaching close to maximal performance with a limited amount of training, but later layers continuing to benefit from increasingly more training. In line with these prior observations, for the Pereira2018 benchmark, we observed that for layers 4-9, performance peaks for the 1M word model, and for the last three layers (layers 10-12), a consistent improvement in performance is observed with larger datasets (**Figure 2E**). This observation echoes prior work showing that later layers build more contextualized representation of linguistic stimuli and better capture syntactic and compositional semantic aspects of the linguistic signal (Belinkov et al., 2017); (Hewitt & Manning, 2019; Tenney et al., 2019), to which the language brain regions are also deeply sensitive (e.g., Fedorenko et al., 2020, 2010; Pallier et al., 2011; Shain et al., 2021).

The general pattern of results was also similar for the secondary, behavioral benchmark (Futrell2018) (**Supp. Figure 3A**): the predictivity of the untrained model and the model trained on 1 million words is significantly below the predictivity of the fully trained model (p-values<0.0001 and p=0.00016, respectively); and the predictivity of the models trained on 10 million words, 100 million words, and 1 billion words does not significantly differ from that of the fully trained model (p-values > 0.05). However, because both of the untrained models achieve reliably above-zero predictivity on the Futrell2018 benchmarks (ps < 0.0001), model performance is unlikely to be related to the representation of linguistic stimuli. As a result, we only present these findings in SI, for completeness.

### 1b. Models that are trained on relatively small amounts of data (a small portion of a massive corpus) can predict human fMRI responses to sentences

In the previous section, we investigated how models that are trained on small corpora (until they reach their best performance on the target language modeling task) perform in predicting human data. However, humans, including children learning a language, are continuously exposed to new words and constructions (see **Supp. Figure 10**). To better simulate such scenarios, as well as to evaluate the robustness of the results to approach, we examined how the ability of a model to predict human fMRI responses to sentences changes over time as the model is being trained on a very large corpus, similar to (Caucheteux & King, 2022). To do so, we used a GPT-2 model that was trained on a corpus consisting of over 9 billion tokens and selected several checkpoints during the training process (0.1%, 1.0%, 10.0%, 100%, and 10 x 100% of training steps, where 100% of training steps approximately equal 1 complete pass over the full dataset; see <u>Methods</u>). At each of these checkpoints, we tested how well the model representations capture human responses to sentences.

For the primary, fMRI (Pereira2018) benchmark, the performance of the fully trained model (i.e., 10 x 100% of training steps) closely matches the results reported in Schrimpf et al. (2021) (where the Hugging Face version of the model was used; cf. the GPT-NEOX library version here), with no significant difference in predictivity (p=0.67). This result shows that model-to-human alignment is robust to the details of model implementation, as one would hope. Critically, mirroring the results from Experiment 1, we observed a consistent increase in how well the model predicts fMRI responses to sentences until the model reaches the 10% checkpoint, at which point the performance plateaus. Critically, the predictivity of the models trained on 10% or 100% of the training steps does not significantly differ from the predictivity of the fully trained model (p-values > 0.05). In contrast, the predictivity of the untrained model (the version with the Hugging Face initialization parameters) and models trained on 0.1% or 1.0% of the training steps is significantly below that of a fully trained model (ps<0.001) (**Figure 2C**; see **Supp. Figure 4** for evidence of robustness to seed choice during model initialization).

The slight decrease in performance with more training (from 100% to 10 x 100%) suggests that more training does not necessarily lead to better alignment with human brain data, although it is possible that this result is due to the relatively spatially and temporally coarse nature of our neural measurements. In particular, a response in a given fMRI voxel reflects an average activity of a large population (a few hundred thousand) of neurons, and the activity is averaged over multiple seconds, which necessarily obscures the fast dynamics of language processing. It is possible that for finer-grained neural data, such as intracranial recordings (electrocorticography or stereo EEG), we might continue to see improvements with more training.

In exploratory analyses of the individual model layers, we observed that performance shows a consistent increase across layers up to the 1.0% checkpoint. After that, the early and middle layers show a drop in performance from the 10% checkpoint to the 10×100% checkpoint, whereas in the later layers, performance increases from the 10% to the 100% checkpoint and then reaches a plateau (**Figure 2F**). Additionally, as in Experiment 1 and in line with prior work in vision (e.g., Geiger et al. 2020; Storrs et al. 2021), earlier layers reach close to maximal performance earlier in the training (at the 1% checkpoint) whereas later layers reach their peak close to the 10% checkpoint (**Figure 2F**).

The pattern of results for the secondary, behavioral benchmark (Futrell2018) closely follows the pattern that we observed with limited-size training datasets in Experiment 1, with predictivity reaching a plateau after the 1% checkpoint (**Supp. Figure 3B**). The predictivity of the untrained model and the models trained on 0.1% or 1% of the training steps is significantly below the predictivity of the fully trained model (p-values: <0.001, <0.001, and 0.0015, respectively); and the predictivity of the models trained on 10%, 100% and 10*100% of the training steps does not significantly differ from that of the fully trained model (p-values > 0.05).

## 2. Model perplexity predicts model performance on the human neural benchmark

For ANN language models, perplexity (a measure of performance on the next-word prediction task; see <u>Analyses</u>) is a reliable predictor of model performance on diverse NLP benchmarks (e.g., Radford et al. 2019; Brown et al. 2020). Schrimpf et al. (2021) further found that off-the-shelf models that perform better on the next-word prediction task are also better able to capture human neural and behavioral responses, in line with prior work showing a similar relationship in pre-transformer models between the amount of training and the ability of a model to predict ERP responses (Aurnhammer & Frank, 2019; S. L. Frank et al., 2015); cf. (Pasquiou et al., 2022).

Here, we examined the relationship between model perplexity and its ability to predict human fMRI responses for models that only differ in the size of the training corpus and for a model at different stages of training, in order to test whether better performance on the next-word prediction task is associated with representations that are more strongly predictive of human neural responses to language.

As expected, perplexity is lower (i.e., the ability to predict upcoming words is better) for models that are trained on larger datasets (**Figure 3A**) and for a given model at the later stages of training (**Figure 3B**; see **Supp. Figure 3C,D** for the results on the behavioral benchmark).

**Figure 3.**
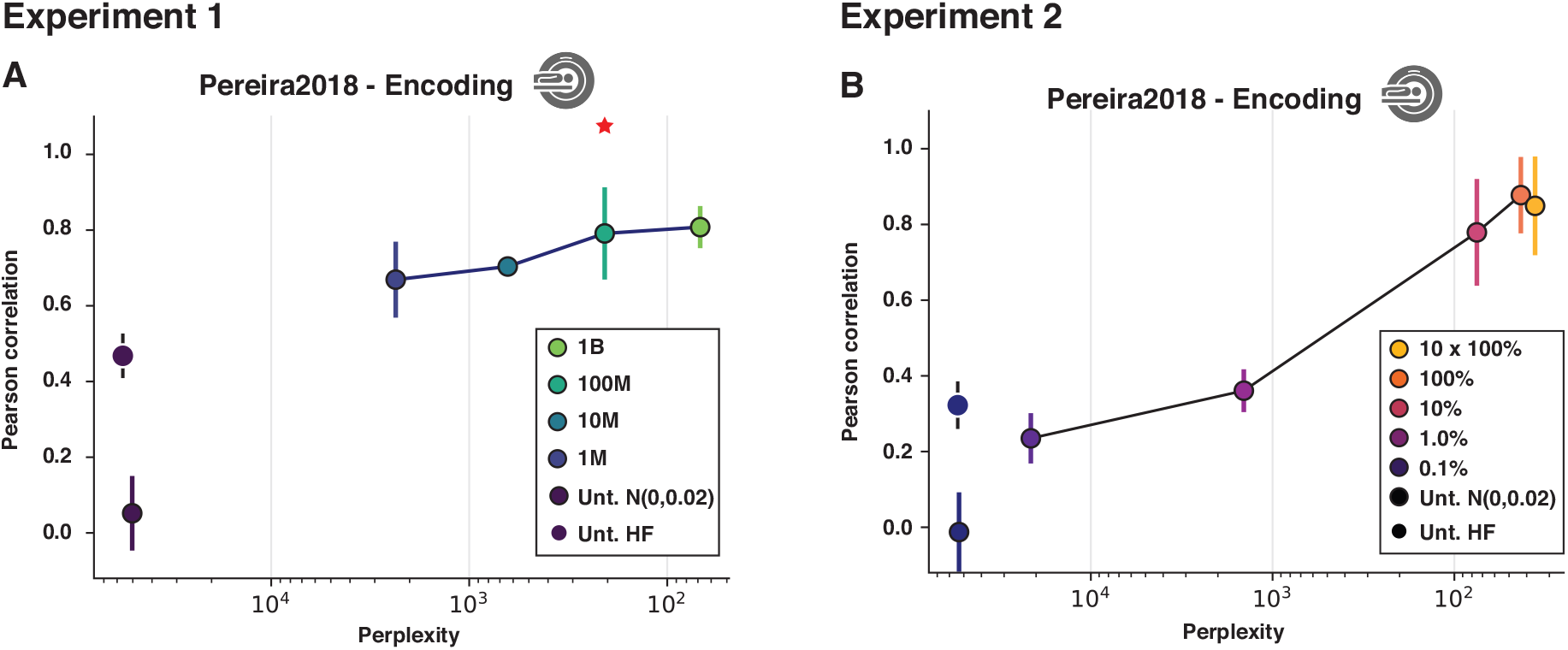
Relationship between model perplexity and model ability to predict human brain responses to sentences. **A.** Experiment 1 results: the relationship between perplexity, i.e., the model’s ability to predict the next token in an independent dataset (wikitext-103-raw-v1), shown on the x-axis, with lower values corresponding to better performance, and model performance in predicting language-responsive voxels’ activation in the Pereira2018 fMRI benchmark. The results are shown for i) two versions of an untrained model (black dots; see Figure 2 caption for details); and ii) four models trained on datasets of different size (1M, 10M, 100M, and 1B words) (blue-to-green dots connected by a line; the model trained on a developmentally plausible amount of data—100M—is marked with a red asterisk). **B.** Experiment 2 results: the relationship between perplexity and model performance in predicting language-responsive voxels’ activation in the Pereira2018 fMRI benchmark. The results are shown for i) two versions of an untrained model (black dots; see Figure 2 caption

Critically, across both Experiments 1 and 2, we observed a consistent relationship between perplexity and neural predictivity, such that lower perplexity is associated with higher predictivity. However, once a model reaches a certain level of perplexity, further improvements in the model’s ability to predict the next word are no longer associated with increases in predictivity, in line with recent findings (Oh & Schuler, 2022, 2023).

## Discussion

In this work, we investigated the relationship between the amount of training data and predictivity of fMRI responses to sentences for transformer-based artificial neural network (ANN) language models. Our study makes several contributions, as summarized next.

### Even when trained on a developmentally realistic amount of data, transformer language models align with human data

Using an fMRI benchmark (Pereira et al., 2018), we established that even with a developmentally realistic amount of training data (∼100 million words, comparable to what humans get during the first 10 years of life; (Hart & Risley, 1992, M. C. Frank, 2023), a GPT-2 model achieves near-maximal predictivity of fMRI responses to sentences. This effect generalizes to a different model architecture (a bidirectional-attention transformer: RoBERTa), although, compared to GPT-2, such models appear to be less sample-efficient, requiring more training data to achieve peak predictivity. (The result also generalizes to a behavioral reading-times benchmark (Futrell et al., 2018), although high performance of untrained models on this benchmark prompted us to move these findings to SI.) In a complementary approach, we showed that when trained on a large dataset, a GPT-2 model already achieves near-maximal predictivity with only 10% of the training steps, well before a full pass over the dataset.

These results align with prior work in vision. For example, Geiger et al. (2020) found that even a small amount of training can result in model representations that are predictive of neural responses in macaques. Moreover, the logarithmic nature of the increase in predictivity between a model trained on 1 million tokens and a model trained on 1 billion tokens aligns with prior Natural Language Processing (NLP) results (e.g., see Kaplan et al., 2020) for evidence of a logarithmic relationship between training data size and the loss in training, and between model size and loss), as well as with vision research (e.g., see Geiger et al. 2020 for evidence of a logarithmic relationship between training data size and predictivity of neural firing rates).

The key implication of these findings is that although large language models are trained on vast amounts of data (and performance on some NLP benchmarks continues to improve with more training), this large amount of training is not necessary for these models to acquire representations that are predictive of human brain responses and behavior. The fact that ANN models trained on a developmentally plausible amount of data can accurately capture fMRI responses to sentences helps address one of the most common criticisms of these models as models of human language processing.

### Alignment between untrained ANN language models and human fMRI responses is strongly affected by the initial unit weight configuration

By relating different versions of untrained models to human fMRI responses, this work clarifies the contributions of architecture to the models’ ability to predict neural responses to linguistic input. Schrimpf et al. (2021) (see also Caucheteux & King, 2022; Pasquiou et al., 2022) have found that untrained models predict fMRI responses quite well, albeit worse than trained models. They speculated that good performance of untrained models might be due to the smoothing of word embeddings across layers in a way that enables the embeddings to capture some aspects of statistical regularities of language (perhaps something as general as nearby words being likely to be related to one another). However, what counts as ‘untrained’ is important to clarify.

‘Untrained’ models come with a particular setting of their unit weights. A particular weight configuration may get ‘baked into’ a model during the process of model development, aimed at maximizing learning efficiency for the target task. Such potential ‘biases’ in initial, pre-trained weights may be akin to innate, evolution-shaped, aspects of brain structure, which may filter information in specific ways as it travels within or across brain areas, even before any learning of the input regularities has occurred (e.g., Zador, 2019). We showed that initializing a model with a normal distribution for all weights leads to the model being unable to predict fMRI response to sentences (predictivity is at ∼0; of course, such a model is also unable to perform the next-word prediction task). (This inability to predict fMRI responses for models initialized with a normal distribution is not due to the lack of activity propagation across layers, as shown in **Supp. Figure 8B**. We also showed that the standard deviation of weight initialization only has minimal effect on the predictivity for untrained model (**Supp. Figure 8D**).

In summary, the ability of untrained models to predict fMRI responses to sentences that was reported in previous studies should not be taken as evidence that model architecture alone (i.e., the units and the patterns of connections among them) can capture human neural responses to linguistic input, or at least, it should be acknowledged that these effects are due to the particular pre-trained *weight configurations*. Furthermore, if a model can (at least partially) match human data with a few bits of information in the form of the initialization parameters (see **Supp. Figure 8C** for evidence that above-baseline predictivity for some initializations may result from the representations for different sentences being more similar), then any results at that alignment level or below for trained models are not meaningful and we should focus on progress beyond that alignment level. Another implication is that future attempts to align trained ANN models with human data should generalize their findings across different weight initializations (Mehrer et al., 2020).

### Model perplexity predicts brain scores

In line with Schrimpf et al.’s (2021) claim that models that perform better on next-word prediction are better at predicting brain data (see also Caucheteux & King, 2022), we found that model perplexity for different amounts of the training data is a good proxy for model performance in predicting responses to sentences. We observed this relationship both in Experiment 1, where we varied the size of the training dataset, and in Experiment 2, where we tested model representations at different points during the training on a large dataset. These findings provide further evidence that optimizing for predictive representations—through training the models on the next word prediction task—may be critical for ANN models to acquire representations that are predictive of human responses to linguistic input.

This finding aligns well with earlier work, which showed that surprisal (how predictable a word is from the preceding context), which is closely related to perplexity, is generally predictive of human behavioral responses (e.g.,Smith & Levy, 2013) and neural responses, as estimated with EEG (e.g., Aurnhammer & Frank, 2019; S. L. Frank et al., 2015; Rabovsky et al., 2018), MEG (Brodbeck et al., 2022; Heilbron et al., 2022), fMRI (Brennan et al., 2016; Heilbron et al., 2022; Henderson et al., 2016; Lopopolo et al., 2017; Shain et al., 2020; Willems et al., 2016), or intracranially (Goldstein et al., 2022) during language processing. However, as recently shown in Tuckute et al. (2023), representations from language models achieve substantially higher predictivity for fMRI response to sentences than more traditional surprisal metrics based on n-gram counts or PCFG parser probabilities.

One recent study (Pasquiou et al., 2022) did not observe a relationship between perplexity and model ability to predict human fMRI responses to linguistic input. We speculate that the lack of this relationship in Pasquiou et al.’s data may relate to the use of an extended-narrative stimulus (the entire ‘The Little Prince’ book) rather than single sentences or short passages. The overall low encoding performance for such stimuli imposes a ceiling on the correlations between model-to-brain alignment and model perplexity (or other variables), making it difficult to differentiate among models. Alternatively, humans and models may use different information for predicting upcoming words, especially in extended linguistic stimuli (Oh & Schuler, 2022).

Why models struggle with predicting neural responses to long narratives is a separate and important question. We offer a speculation. In the human brain, division of labor exists between i) the language-selective network, which integrates information within clauses/sentences but does not track longer-range contexts (e.g., Blank & Fedorenko, 2020), and ii) the Default network(s) (Buckner & DiNicola, 2019), which integrates information over extended temporal contexts (Lerner et al., 2011). Importantly, the Default network does not operate over word sequences; instead, the information that this system represents is likely abstract, as evidenced by the fact that it processes long contexts in both linguistic and non-linguistic stimuli (e.g., Baldassano et al., 2017; Simony et al., 2016). As a result, the ANN language models (like those used in current work and in (Pasquiou et al., 2022) may simply lack representations that are sufficiently abstract (not directly tied to the stimulus, i.e., to the word sequences) to match those in the Default network, perhaps because language models eventually have to ‘go back’ to specific words in order to perform the next-word prediction task. Some of the newer models, like GPT-3, seem to be able to handle a greater degree of abstraction (Brown et al., 2020) and thus may be promising for future attempts to capture human neural responses to long and complex linguistic stimuli.

### Limitations and future directions

In general, it is challenging to compare the amount of training data that a model gets to the amount of linguistic input that a child gets. The consequences of a single token of input for a computational model depend on many aspects of the model’s architecture, training setup, and so on; and of course, the fact that, for smaller datasets, the same dataset is repeated multiple times during the training varies drastically from what humans experience. All the results should therefore be interpreted in light of these limitations.

Furthermore, we have here focused on the effects of the *amount* of training data on the ANN language models’ ability to capture human responses to language. However, the *nature* of the training data is, no doubt, also important. For example, training models on data that are similar to what children are exposed to could lead to improved neural predictivity (Chang & Bergen, 2021; Warstadt & Bowman, 2019). Indeed, this approach has been shown to improve vision models’ ability to capture primate neural responses (Mehrer et al., 2021). It will also be important to investigate the role of the *learning algorithms* that the models use and their *training objective*, as both likely affect the representations that the models learn (e.g., see Zhuang et al., 2022 for evidence from vision). Specifically, Zhuang et al. (2022) showed that in an object categorization task, the negative sampling objective function, which maximizes the similarity between objects in the same category while minimizing the similarity between objects in different categories in the internal representation of the model, can alleviate model failure in capturing human visual behavior, which occurs under the standard objective function. This failure is due to the presence of categories that are infrequent in the training data, and this finding can be relevant for language, which also contains infrequent elements (words and constructions) amid more common ones.

Another issue that we did not investigate here is the nature of various training parameters (e.g., learning rate, batch size, randomization of context, length of context, etc.). Such parameters can affect model performance on NLP tasks and, possibly, their ability to predict human neural or behavioral responses to language. However, we suspect that the influence of these parameters would be relatively minimal given the evidence from prior work that model size, dataset size, and amount of computing power are the main contributors to model performance after training, as measured in loss for predicting the next token (Kaplan et al., 2020), and that model size is the main contributor to model performance in predicting fMRI responses to language (Antonello et al., 2023).

Another aspect of the ANN models that is important for building accurate models of human language processing is the model *architecture*. We here generalized our training effects across uni- and bi-directional-attention transformers, but a systematic investigation of the effects of diverse architectural parameters (e.g., the number and size of layers, number of attention heads, etc.) on the models’ ability to predict human responses to language would be valuable.

Tightly controlled comparisons between different classes of model architectures are more challenging but creating numerous model variants all trained on the same dataset (e.g., Storrs et al., 2021) could enable identification of architectural motifs that are essential for a good match with human neural and behavioral data.

Perhaps the biggest limitation of this and related work is the obscurity of both the model representations and human neural representations. It is not known what aspects of the representations change as the models are trained on increasingly more data (aside from knowing that these changes lead to improved performance on the next-word prediction task), and how exactly these changes in the models’ representations of linguistic input impact their ability to predict brain activation or behavioral processing difficulty. Some recent work has begun to attempt isolating the aspects of model representations that affect model-to-brain alignment. For example, (Kauf et al., 2023 ; current special issue) performed a series of experiments where model representations were obtained for different perturbations of a linguistic stimulus (e.g., scrambling the word order or dropping/replacing some of the words) and then related to neural representations of an intact stimulus in order to see which perturbations affect model representations negatively. They found that word-level and compositional semantic information appears to be more important than information related to the syntactic structure in the model-to-brain alignment. Still however, much about the details of how models vs. humans represent and process linguistic stimuli remains to be discovered.

Another exciting recent approach (Sexton & Love, 2022) is to replace a part of a model (the one best aligned with human neural responses) with fMRI signals to test whether parts of the model representation that align with neural data affect model behavior.

In future work, we aim to address these gaps in order to build increasingly more accurate and interpretable models of language processing in the brain.

## Acknowledgements

This work was partially supported by NIH award U01-NS121471 to EF. EAH and MS were supported by the Friends of McGovern graduate fellowships. NZ was supported by a K. Lisa Young ICoN Center postdoctoral fellowship. EF was additionally supported by NIH awards R01-DC016607 and R01-DC016950, as well as by research funds from the McGovern Institute for Brain Research, the Brain and Cognitive Sciences department, the Simons Center for the Social Brain, and the Middleton Professorship. EAH is grateful to Josh McDermott and members of the Fedorenko Lab (especially Carina Kauf, Cory Shain, Greta Tuckute, and Chengxu Zhuang) for helpful discussions and comments on the drafts of the manuscript. EAH is also grateful to Stella Biderman, EleutherAI, with setup in experiment 1, Jason Bolton, Laurel Orr, and Siddharth Karamcheti for help with setup in experiment 2.

## Contributions

EAH, NZ and EF designed research, EAH performed research, MS, YZ, and SB contributed analytic tools, EAH analyzed data with supervision from EF, EAH and EF wrote the paper with input from MS, YZ, SB, and NZ.

## Footnotes

1 https://huggingface.co/nyu-mll

---

**Supplementary Figure 1.**
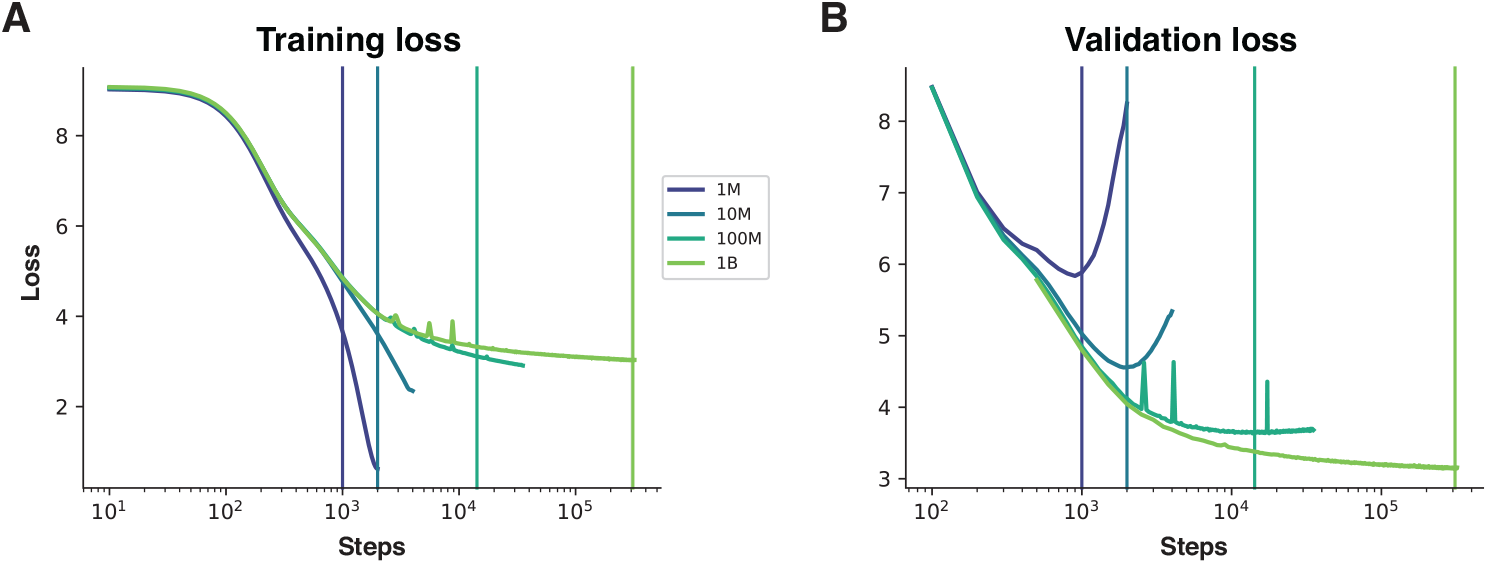
Training and validation loss for the models in Experiment 1. For all models, a training batch size of 64 was used (each element in a batch is a sequence of 1,024 tokens). The training was done with a standard Adam optimizer (LR: 0.0006, beta: [0.9,0.999], eps:1e-8), and cosine learning rate (starting at 0.0006 and decaying over 320000 training steps), with gradient accumulation and clipping, attention and hidden dropout ratio of 0.1, and no weight decay. **A.** Training loss: as expected, all models show a drop in loss with additional training, and for models trained on smaller datasets, the rate of learning is faster. **B.** Validation loss: as expected, all models show a drop initially, and models trained on smaller datasets reach their minimum loss earlier in the training process and have higher values (corresponding to worse performance) than models trained on larger datasets. In both A and B, the vertical lines mark the step with the minimum validation loss (best model performance). The representations for the steps shown in B were used for testing against the human benchmarks.

**Supplementary Figure 2.**
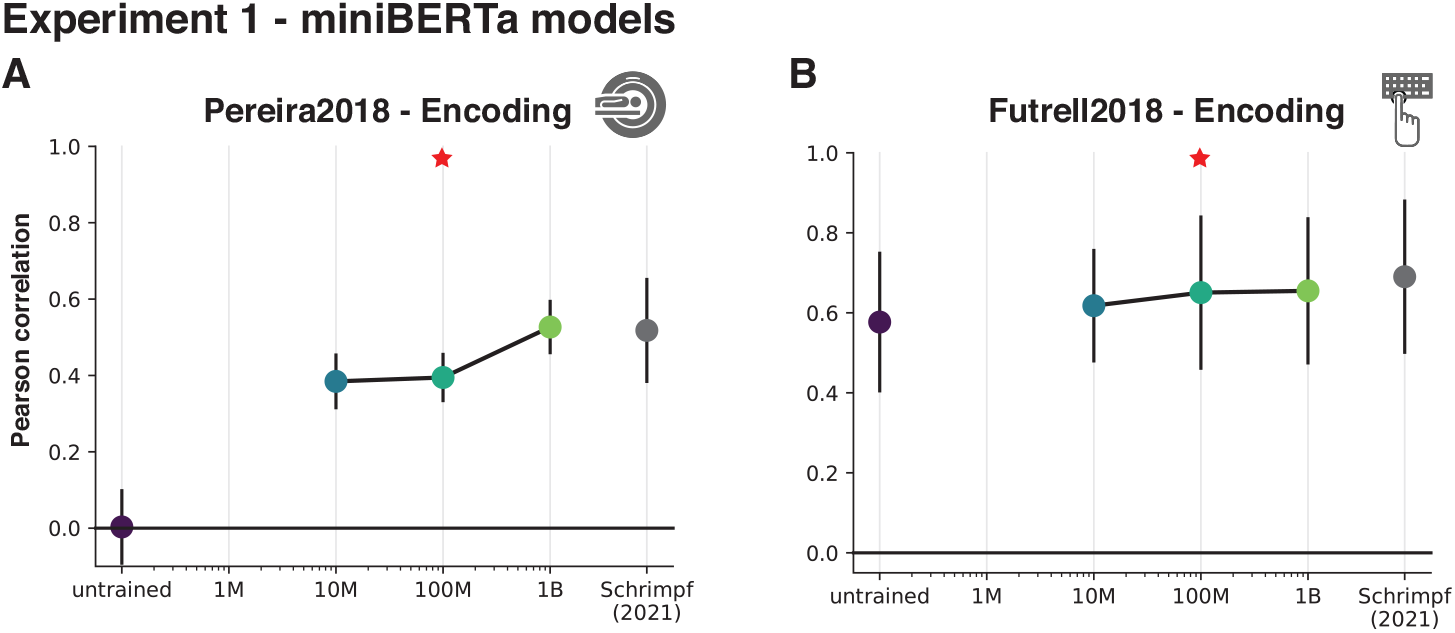
Model performance on the human benchmarks for the miniBERTa models as a function of training in Experiment 1. Performance (normalized predictivity) of the best-performing layer, as reported in Schrimpf et al. (2021) for the RoBERTa model, in predicting language-responsive voxels’ activation in the Pereira2018 fMRI benchmark (A), and reading times in the Futrell2018 behavioral benchmark (B) (see **Supp. Figure 3** for the results for GPT-2 for the Futrell2018 benchmark). The results are shown for i) an untrained model (black dots; the Gaussian initialization; see <u>Methods</u> for details); ii) three models trained on datasets of different size (10M, 100M, and 1B words) (teal-to-green dots connected by a line; the model trained on a developmentally plausible amount of data—100M—is marked with a red asterisk); and iii) a fully trained model, as reported in Schrimpf et al. (2021) (grey dots). As in the main figures, we computed a median score across participants and divided it by an estimated ceiling value to get a normalized score, and we computed a median absolute deviation over participants for use as error bars.

**Supplementary Figure 3.**
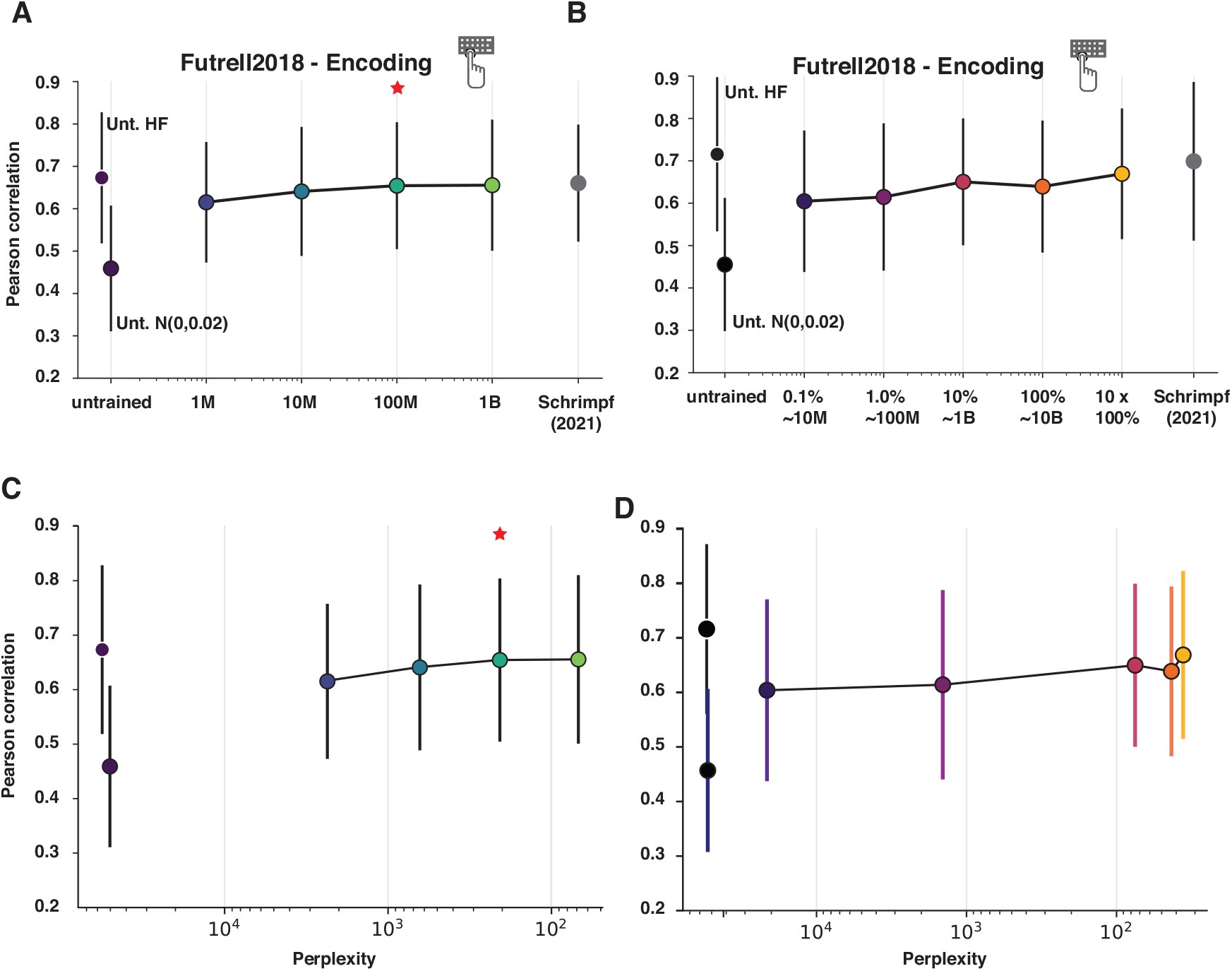
Model performance on the behavioral (Futrell2018) benchmark as a function of training. **A.** Performance (normalized predictivity) of the best-performing GPT-2 layer, as reported in Schrimpf et.al. (2021), in predicting reading times in the Futrell2018 benchmark. The results are shown for i) two versions of an untrained (Unt.) model (initialized in two different ways: Unt. N(0,0.02) corresponds to the untrained model initialized with a mean of 0 and a standard deviation of 0.02, and Unt. HF corresponds to the untrained model initialized with the Hugging Face parameters; see <u>Methods</u>) (black dots); ii) four models trained on datasets of different sizes (1M, 10M, 100M, and 1B tokens) (blue-to-green dots connected by a line; the model trained on a developmentally plausible amount of data—100M—is marked with a red asterisk); and iii) a fully trained model, as reported in Schrimpf et al. (2021) (grey dots). As in the main figures, we computed a median score across participants and divided it by an estimated ceiling value to get a normalized score, and we computed a median absolute deviation over participants for use as error bars. **B.** Performance of the last GPT-2 layer in predicting reading times in the Futrell2018 benchmark. The results are shown for i) two versions of an untrained model (initialized in two different ways, as in A; see <u>Methods</u>) (black dots); ii) a model trained on a large dataset examined at different points during the training (0.1%, 1.0%, 10%, 100%, and 10*100% of training steps) (purple-to-yellow dots connected by a line); and iii) a fully trained model, as reported in Schrimpf et al. (2021) (grey dots). **C.** The relationship between perplexity, i.e., the model’s ability to predict the next token in an independent dataset (wikitext-103-raw-v1), shown on the x-axis, with lower values corresponding to better performance, and model performance in predicting human reding times in the Futrell2018 benchmark. The results are shown for i) two versions of an untrained model (initialized in two different ways, as in A; see <u>Methods</u>) (black dots); and ii) four models trained on datasets of different sizes (1M, 10M, 100M, and 1B words) (blue-to-green dots connected by a line; the model trained on a developmentally plausible amount of data— 100M—is marked with a red asterisk). **D.** The relationship between perplexity and model performance in predicting reading times in the Pereira2018 benchmark. The results are shown for i) two versions of an untrained model (initialized in two different ways, as in A; see <u>Methods</u>) (black dots; see Figure 2 caption for details); and ii) a model trained on a large dataset examined at different points during the training (0.1%, 1.0%, 10%, 100%, and 10*100% of training steps) (purple-to-yellow dots connected by a line).

**Supplementary Figure 4.**
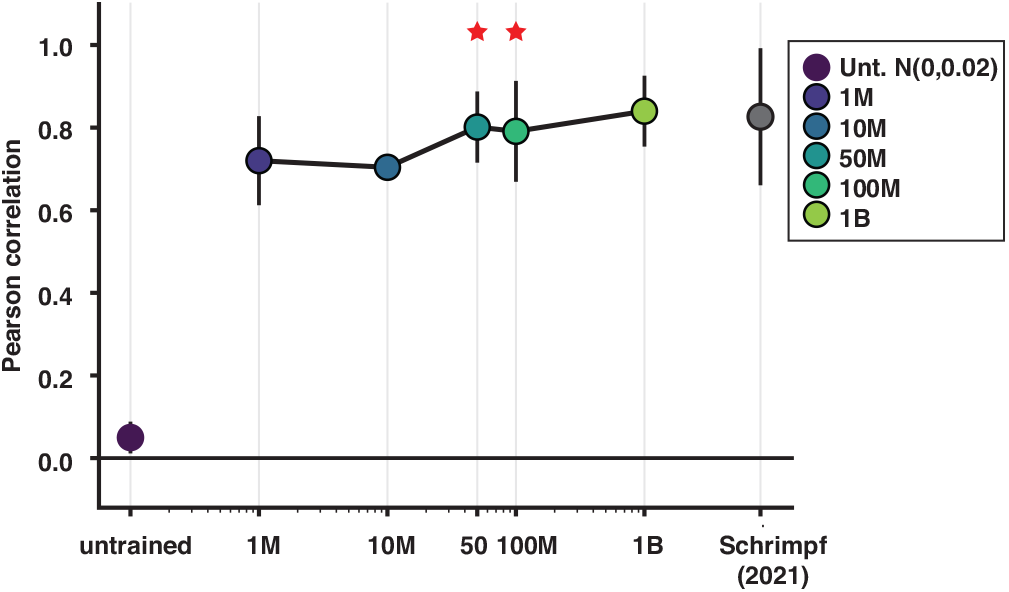
Model performance on the fMRI (Pereira2018) benchmark as a function of training, including a model trained on 50M words (the data for the models other than the 50M model are identical to Figure 2A). Performance (normalized predictivity) of the best-performing GPT-2 layer, as reported in Schrimpf et.al. (2021), in predicting language-responsive voxels’ activation in the Pereira2018 fMRI benchmark. The results are shown for i) an untrained model (initialized with a Gaussian; see <u>Methods</u>); ii) five models trained on datasets of different size (1M, 10M, 50M, 100M, and 1B words) (blue-to-green dots connected by a line; the models trained on developmentally plausible amounts of data—50M and 100M—are marked with a red asterisk); and iii) a fully trained model, as reported in Schrimpf et al. (2021) (grey dots). As in Figure 2, we computed a median score across participants and divided it by an estimated ceiling value to get a normalized score, and we computed a median absolute deviation over participants for use as error bars. The models trained on 50M, 100M and 1B words exhibit fMRI response predictivity that is similar to that of the fully trained GPT-2 model, with no significant differences in predictivity values (ps>0.05). In contrast, the predictivity of the untrained model and the models trained on 1M and 10M words is significantly below the predictivity of the fully trained model (p-values: <0.0001, p=0.001, and p=0.003, respectively).

**Supplementary Figure 5.**
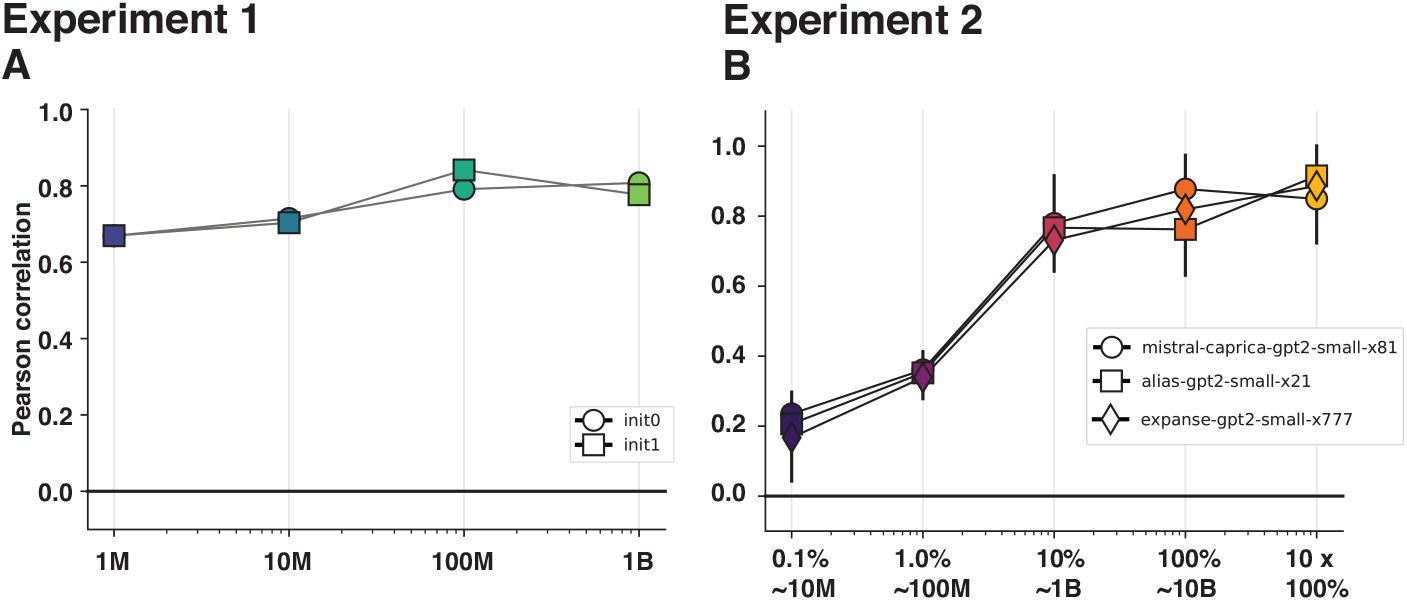
Robustness of the critical results to seed choice during model initialization. Model initialization seeds have been shown to affect model performance (e.g., Mehrer et al., 2020). Here, we show that the main results (reported in Figure 2A,C) are robust the seed choice. For Experiment 1, two different seeds were used; for Experiment 2, three different seeds were used.

**Supplementary Figure 6.**
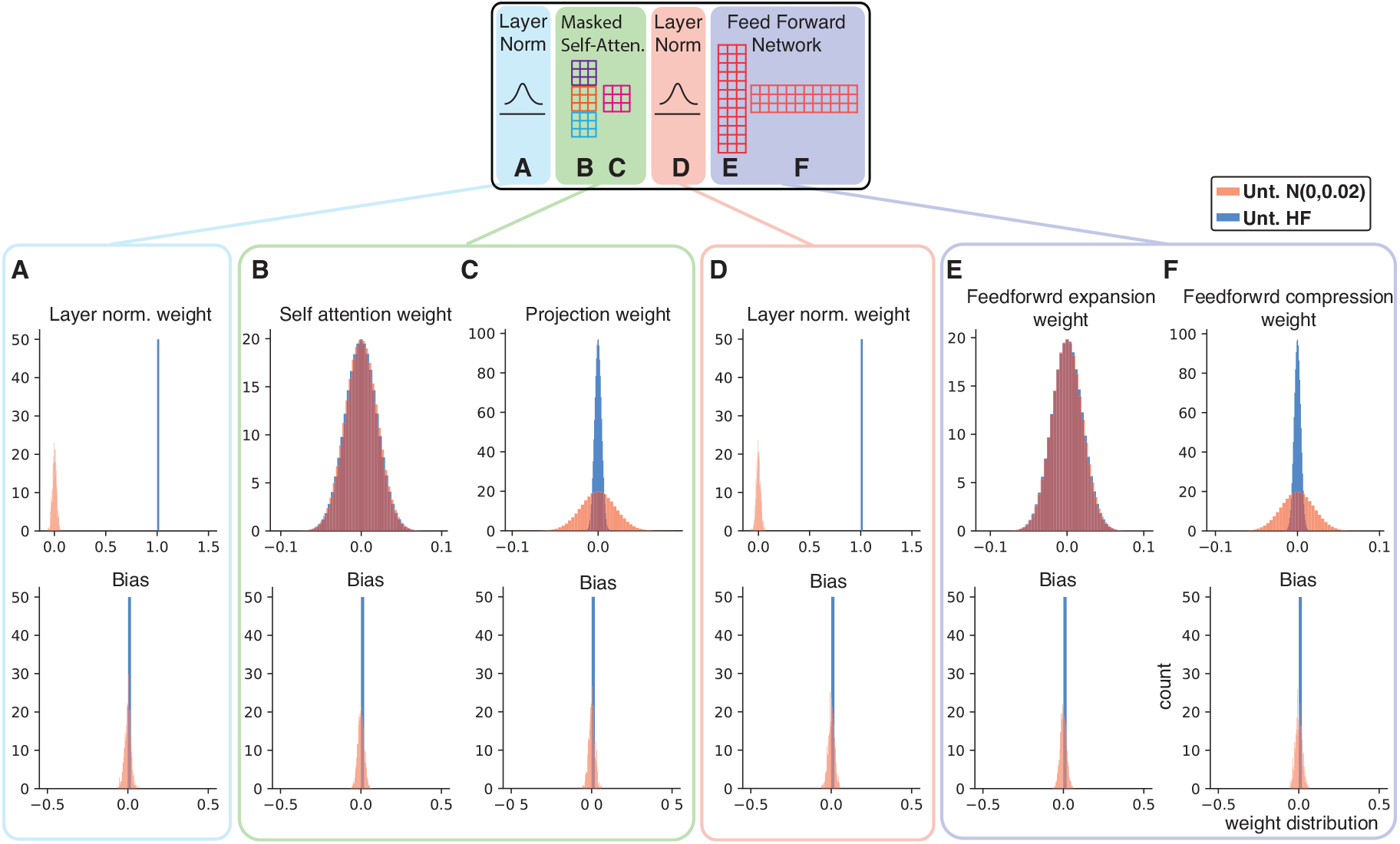
A detailed comparison of the two model initializations for the untrained GPT-2 models (for a sample layer (layer 1)). The distribution of weight (top row) and bias (bottom row) values for the first layer normalization (A), two operations in self attention (B-C), the second layer normalization (D), two operations in feedforward processing (E-F) for two versions of an untrained model: Unt. N(0,0.02) corresponds to the untrained model initialized with a mean of 0 and a standard deviation of 0.02, and Unt. HF corresponds to the untrained model initialized with the Hugging Face parameters (see <u>Methods</u>).

**Supplementary Figure 7.**
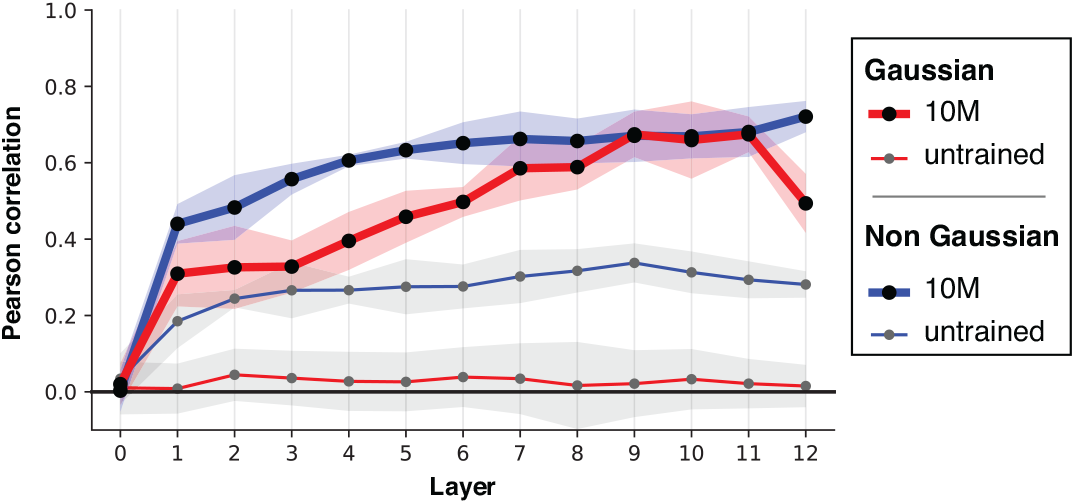
Effects of differences in model initialization on the performance of an untrained GPT-2 model and a GPT-2 model trained on the 10M words on the Pereira2018 fMRI benchmark across layers. Performance (normalized predictivity) in predicting the Pereira2018 fMRI benchmark. The models shown with blue lines were initialized with the Hugging Face parameters; the models shown with red lines were initialized with a Gaussian distribution of weights with a mean of 0 and a standard deviation of 0.02 (see <u>Methods</u> and **Supp. Figure 6**). The untrained model initialized with a Gaussian distribution performs close to 0 across layers. In contrast, the untrained model initialized with the Hugging Face (non-gaussian) parameters achieves ∼0.4 predictivity for some layers. After training, both models reach a similar level of predictivity for their later layers (layers 9-11).

**Supplementary Figure 8.**
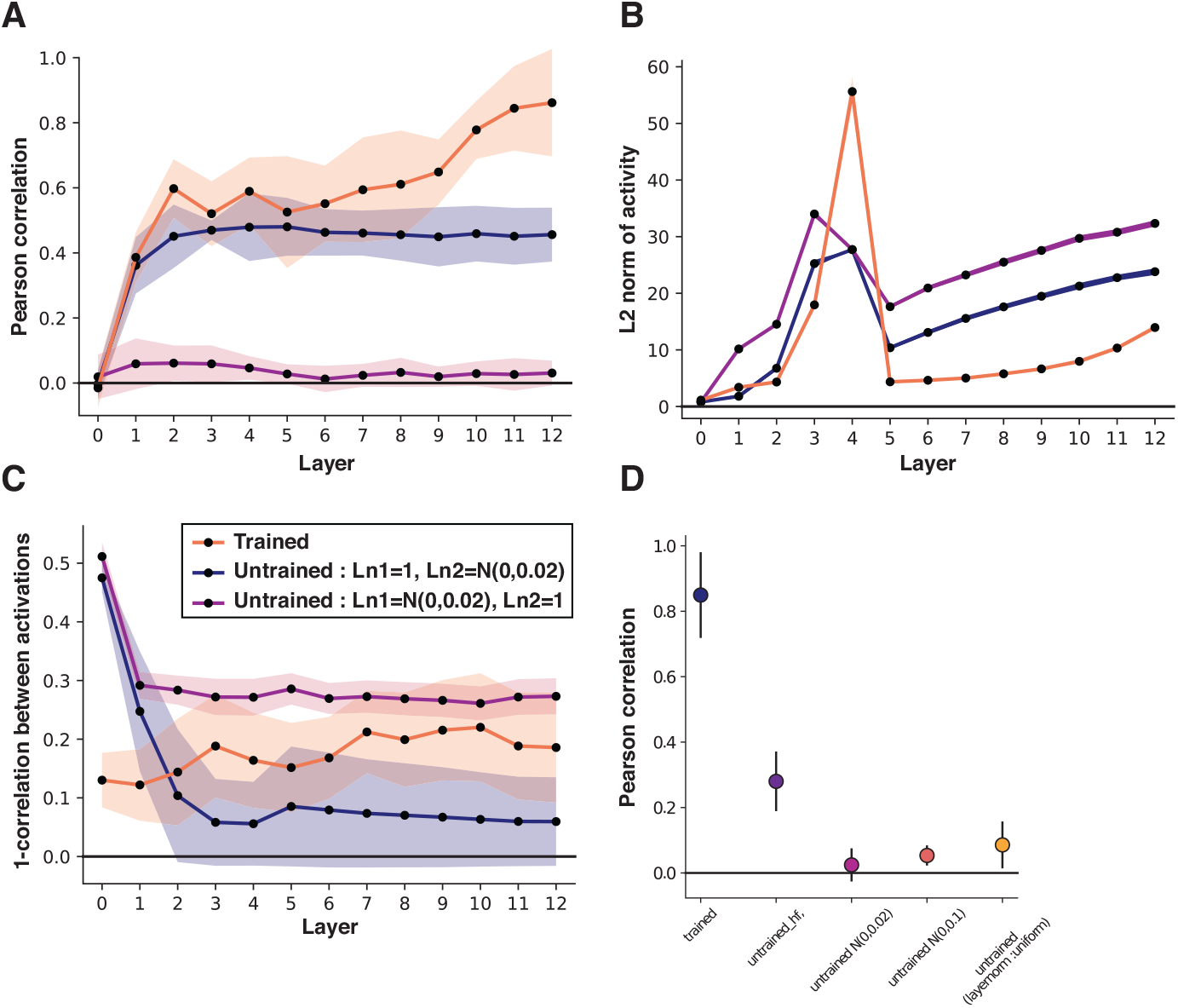
Effects of differences in model initialization on the performance of untrained models on the Pereira2018 benchmark across layers. Here, we created two versions of an untrained model that both use a Gaussian weight distribution for the self-attention and feedforward processing components of the model but differ in the weights they use for layer normalization. In particular, in one model (the dark blue line), the first layer normalization (Ln1) was set to 1 (as in the Hugging Face initialization), and the second layer normalization (Ln2) was set to a Gaussian distribution, and in the other model (the maroon line), Ln1 was set to a Gaussian distribution, and Ln2 – to 1 (as in the HF initialization). The third line (orange) corresponds to a trained model (initialized with the standard HF parameters) and is included here for comparison. **A.** Performance (normalized predictivity) in predicting the Pereira2018 fMRI benchmark. The model with Ln1=1 exhibits higher performance compared to the model with Ln2=1, which suggests that the first layer normalization plays a larger role in contributing to above-zero performance for untrained models on the human benchmarks. **B.** Amplitude of model activation. The two untrained models show a similar level of activation to each other (and, in some layers, to the trained model), which suggests that the difference in performance between them is not due to the lack of activity propagation across layers in the model where Ln2=1. **C.** Similarity of model representations among the sentences in the Pereira2018 benchmark (higher values correspond to lower similarity). The representations appear to be more similar in the model where Ln1=1 (compared to the model where Ln2=1 or compared to a trained model), which suggests that setting Ln1 to 1 effectively removes stimulus-specific encoding and may explain the above-zero predictivity of neural responses for the untrained model initialized with the HF parameters. **D.** Performance of a trained model and several versions of untrained models in predicting the Pereira2018 fMRI benchmark. The untrained models include a model initialized with the standard HF parameters, two models initialized with a Gaussian distribution that vary in the size of the standard deviation (0.02, 0.1), and a model where all LayerNorm weights are set to be a uniform distribution between 0 and 1 (all positive values). The model initialized with the HF parameters shows higher predictivity than the other untrained models, but the size of the standard deviation in the Gaussian initialization does not strongly affect performance (cf. (Rohde & Plaut, 1999)). Finally, the model initialized with a uniform distribution (of positive values) for LayerNorm weights performs close to zero, similarly to the model with a Gaussian initialization.

**Supplementary Figure 9.**
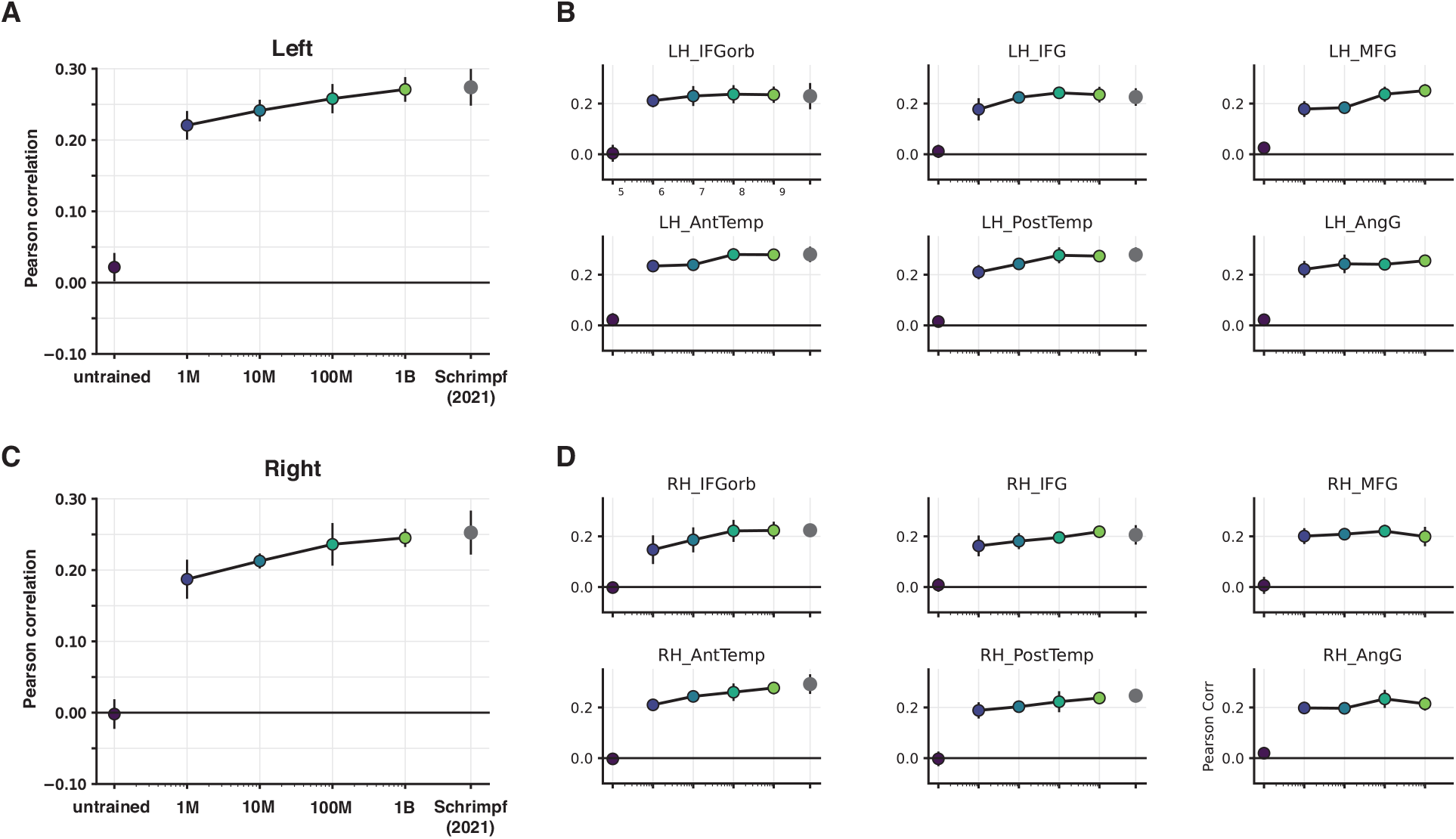
Model performance on the fMRI (Pereira2018) benchmark as a function of training in the left vs. right hemisphere separately (A and C), and in the individual regions of interest within each hemisphere (B and D). Performance of the best-performing GPT-2 layer, as reported in Schrimpf et.al. (2021), in predicting language-responsive voxels’ activation in the Pereira2018 fMRI benchmark for the left hemisphere (A) and right hemisphere (C) overall and broken down by functional ROI (B: left hemisphere (LH) fROIs; D: right hemisphere (RH) fROIs). Note that we here report *raw* predictivity values (cf. the normalized values reported in all other figures) so as to be able to more meaningfully compare between hemispheres and among ROIs without the complication of different ceiling values across regions. The fROIs are identified with an independent localizer, as described in <u>Methods</u>. The general pattern of results (presented in main Figure 2) holds across hemispheres (although predictivity is higher in the LH, in line with other work; e.g., Schrimpf et al., 2021; Tuckute et al., 2023) and fROIs.

**Supplementary Figure 10.**
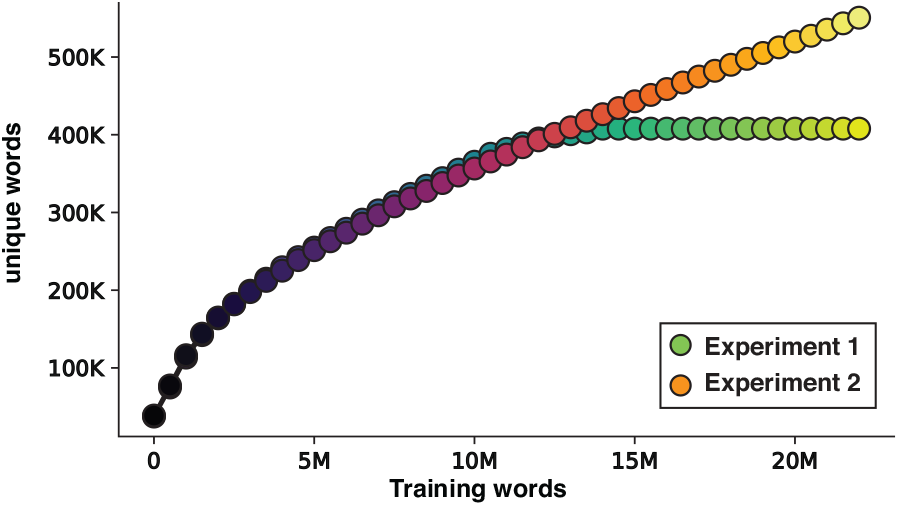
Comparison of the number of unique words during the course of training in Experiment 1 vs. Experiment 2. To illustrate the difference between the setup in Experiment 1 vs. 2, we examined the number of unique words relative to the total number of words seen for the first 20M words, in bins of 500K words. For Experiment 1, we focused on the 10M dataset. As can be seen, the green-to-yellow curve reaches a plateau at 10M words, as the model is no longer exposed to new words after that point; instead, the same 10M training dataset is presented again until the best model perplexity is reached. In contrast, the purple-to-yellow curve continues to increase beyond the 10M mark as the model is continually exposed to new words during the course of training (of course, eventually, the increase in new words will become very small because encountering new words becomes less and less likely after exposure to a large enough training corpus).

